# Recent immigrants from islands dominate two keystone forest herbivore assemblages in Australia

**DOI:** 10.64898/2026.10.13.758981

**Authors:** Paul M. Oliver, Andrew F. Hugall, Flavien Ferreira

**Affiliations:** Centre for Planetary Health and Food Security, Griffith University, 170 Kessels Road, Brisbane, Queensland, 4111, Australia; Biodiversity and Geosciences, Queensland Museum, South Brisbane, Queensland, 4101, Australia; Department of Sciences, Museums Victoria, Melbourne, VIC 3001, Australia

**Keywords:** island biogeography, megabats, Melanesia, pigeons, tropical rainforest, upstream colonisation

## Abstract

Successful upstream colonisation from islands to continents has now been documented in many taxa, however as yet there are no examples of upstream dispersal playing a dominant role in the assembly of regional biotas. Here we use supermatrix phylogenies for two groups of highly mobile volant keystone herbivores – nocturnal megabats and diurnal pigeons – to show that Australian rainforest communities in both groups are entirely derived from at least 17 independent colonisations from the islands of Melanesia and Wallacea. These colonisations are concentrated within the last 5 million years and there is also no evidence of *in situ* diversification post colonisation of Australia. This inference contrasts against perceptions of Australia’s rainforests being dominated by ancient Gondwanan relicts. Furthermore, given the importance of megabats and pigeons for pollination and fruit dispersal in forest environments, these data indicate that recent upstream colonisation from the islands of Melanesia through the late Tertiary likely had a major impact on the floral composition and ecological functioning of Australia’s lowland forests.

## 1. Introduction

A prediction to emerge from island biogeography theory was that successful ‘upstream’ biotic dispersal from islands into larger and more biotically complex systems should be relatively rarer than ‘downstream’ dispersals in the converse direction [1]. This prediction was underpinned by assumptions that larger landmasses will both have more potential propagules, and also be characterised by more “competitive” biotic communities less permeable to new colonists [1,2]. Counter to this prediction, over the last 30 years a range of studies have provided evidence that upstream dispersals from islands to continents have occurred in many taxa [3], and have even led to extensive speciation and diversification [4]. However, analyses of biotic assembly and dispersal at regional scales still tend to indicate that downstream colonisations from continents to islands are more pervasive [5,6]. As yet, there are no examples of geographically or temporally concordant periods of upstream colonisation dominating over downstream colonisation as the source of complete assemblages or regional biotas.

The Australasian region comprises Australia (continental) and many nearby islands to the north (centred on New Guinea, and hereafter referred to as Melanesia). New Guinea is the largest, highest and for many taxa most biodiverse island in the world [7,8]. Biotic exchange between neighbouring Australia and New Guinea has been extensive, and may have played a key role in the assembly of respective biotas [9]. The juxtaposition of a relatively small and climatically unstable continent [10] against a megadiverse assemblage of islands, might potentially predispose these two regions to showing a strong signature of upstream colonisation. Evidence for this prediction is mixed. For some radiations such as marsupials [11], passerine birds [12] and various taxa of plants [13], a deep history in Australia and initial dispersal out of Australia is supported, with subsequent diversification in other parts of the world. In contrast, rats [9], some lineages of birds [14–16] and plants [17] all show signals of upstream colonisation of Australia from tropical regions of Melanesia. In yet other groups, rates and numbers of dispersal between the two regions are very low, suggesting the existence of major environmental barriers to dispersal between the two regions [18].

Broadscale analyses of the origins of the Australian rainforest plant fauna have particularly emphasised that the tropical lowland or “megathermal” forests of Australia have a high proportion of species with close relatives in regions to the north [17,19]. To date, many of these studies have not differentiated between biotas of Asia and those of Melanesia, and have thus not explicitly considered a role for upstream colonisation in shaping this recent influx of lineages into megathermal environments. However, a recent study of pigeon biogeography and dispersal found that Australian rainforest assemblages of fruit-eating species were dominated by recent Plio-Pleistocene immigrants from Melanesia [15].

Here we present a synthesis of the biogeographic origins of Australian forest-dwelling pigeons and one other lineage of herbivores in tropical Australia – the Megachiroptera, commonly referred to as fruit bats or megabats. Both lineages are considered keystone fruit and pollen dispersers in tropical forest ecosystems [20,21]. Using the most species-complete phylogenetic datasets for megabats and pigeons assembled we here focus on resolving: i) patterns of lineage accumulation and endemic diversification in both Australia and nearby insular systems; and ii) the number, timeframe and directions of dispersal events involving Australia. These data show that Australian rainforest communities in both radiations are dominated by taxa derived from at least 17 independent upstream colonisations, of which the vast majority occurred recently, and also show negligible signatures of subsequent endemic diversification post colonisation of Australia. These data provide a novel demonstration of ecologically comparable species assemblages wherein recent upstream colonisation dominates as the source of a continental biotic assemblage.

## 2. Methods

### 2.1. Taxonomic and Distribution Information

Taxonomy, nomenclature and distribution information for megabats were derived from OZCAM (https://ozcam.org.au/)[22] and *Bats of the world* (https://batnames.org/query.html)[23], and for pigeons from the International Ornithology Council World Bird List v. 9.2 (https://www.worldbirdnames.org/updates/). Additional information for both taxa was taken from Wikipedia, the IUCN redlist (https://www.iucnredlist.org/) and relevant phylogenetic literature (Supplementary file Appendices). For the megabats we further rationalized sequences to additional subspecies/geographic units/distinct genetic units based on recognized nomenclature and genetic data (summarised in Supplementary files).

### 2.2. Supermatrix Assembly

We built our phylogenetic supermatrix databases based on three key components: i) multi-gene sequence data downloaded from the NCBI global sequence; ii) a phylogenomic datasets to provide a strong backbone across the major lineages, both in topology and relative branch length; and iii) nomenclature reconciled to a taxonomic database. Collation and curation of multiple data sources followed supermatrix assembly procedures used previously [15,24] including custom Bash scripts.

For the Megabats we incorporated 10 genes widely used in previous Pteropodidae supermatrix work (e.g., Almeida et al. [25,26], Giannini and Simmons [27]) with additional phylogenomic data [28]. For Pigeons we updated the dataset of Oliver et al. [15] with the new phylogenomic backbone of Oswald et al. [29] and new NCBI data (https://www.ncbi.nlm.nih.gov).

#### 2.2.1. Multi-gene sequence data

Data were downloaded from the NCBI nucleotide collection. We used gene sequences obtained for key phylogenetic references through BLAST+organism searches with an E-value threshold of 5e-6 and retained aligned regions. For protein coding genes and ribosomal RNA genes we used tblastn and blastn respectively. Gene sequences were aligned with MAFFT v. 7.505 [30] using the ‘AUTO’ setting. We then assembled alignments into a custom Microsoft Excel database, and reconciled nomenclature. We inferred gene trees through IQ-TREE 2.4.0 [31] using allnni and models of sequence evolution as identified by ModelFinder implemented in IQ-TREE [32].

Rather than selecting one ‘best’ longest species sequence per gene, we retained all sequence data to first assess the gene tree monophyly of nominate species as well as genera [24], revising the dataset by removing anomalous sequences and in some cases updating identifications. Some genes were extracted from genome assemblies, mitogenome accessions and SRA files. We also included mtDNA data that came from Nesi et al. [28] megabat phylogenomic study.

#### 2.2.2. Higher-level phylogenomic backbone

The recent phylogenomic work provided the key backbone for the Megabats [28], spanning 114 species in 43 genera (Alignment_Concat_Nuc1455_complete.nex https://doi.org/10.5061/dryad.7m0cfxpq5). This 2,791,426 site dataset was first trimmed of sites available for less than 95% of taxa, then jack-knifed while stochastically depleting large blocks (>20) of invariant sites, down to a final length of 30,840 sites 98.2% complete. The same process was implemented for a published Pigeon phylogenomic matrix of 73 species spanning 38 genera (AlignmentMatrix.phy doi:10.6084/m9.figshare.c.7589910)[29], reducing it from 10,586,851 sites to 31,437 sites that were 99.4% complete. Trees were then inferred from these phylogenomic ‘stubs’ using IQTree [33](settings TN93+F+G4 model with-allnni and-bnni 1000 ultrafast bootstrap support). Resultant trees (Figure S1) are reasonable representations of the original all-data trees (albeit with reduced support on some nodes).

#### 2.2.3. Supermatrix assembly and analysis

Concatenated supermatrix sequence data used the best (longest accepted) single exemplar sequence per gene per species, for all species with a total of >500 bases of any data. These gene alignments were then compacted by removing regions with little or no data (<10% taxa per gene). Sparse, patchy supermatrices often include taxa groups that share no data in common, which can bias phylogenetic inference [34,35]. We used the simple measure of the number of genes taxa have in common (GIC) [24], to assess supermatrix data overlap. For the Megabats we used outgroups from Nesi et al. [28], augmented with some NCBI multi-gene data. For pigeons we updated our composite higher taxa [14] with the passerine phylogenomic dataset of Oswald et al. [29] assigned to the ‘neoaves’ outgroup.

Table S1 and S2 provide a complete breakdown of taxa, genes, sites, and sources of gene data (e.g. NCBI accessions, phylogenomic data) for both the Megabat and Pigeon supermatricies. Tables S3 and S4 summarize genes used and sites in the final Megabat and Pigeon supermatrices. Here the term ‘gene’ refers to distinct genetic data unit. The combination of the phylogenomic backbone frameworks and the standard mitochondrial ‘barcode’ genes (cytb, ND2, COI) provide considerable data overlap among taxa in both supermatrices.

The final megabat supermatrix includes 204 ingroup taxa and 6 outgroups built from 1,260 data units in 11 genes, comprised 4,637,265 bases in 42,871 sites (38,058 nuclear, 4,813 mitochondrial), with median values of 77% site-complete, 32,894 bases per taxon and 6 genes per taxon. Pairwise GIC indicated extensive overlap across all ingroup taxa (zero for 2.3% of taxa, one for 16%) and within genera (zero 0.7%, one 13%).

The Pigeon supermatrix comprises 252 ingroup taxa and 3 outgroups built from 1,476 data units in 12 genes, comprised 3,477,546 bases in 44,150 sites (36,759 nuclear, 7,391 mitochondrial), with median values of 12% site-complete, 5,118 bases per taxon and 5 ‘genes’ per taxon. Pairwise GIC were generally good across all ingroup taxa (zero for just 3.7%, one 23%) and within genera (zero 0%, one 10%).

### 2.3. Phylogenetic inference

Trees were inferred from the supermatrix alignments using RAxNG v1.2.2 [36] and IQ-TREE 2.4.0, with optimized partition model of sequence evolution (Tables S3 and S4) identified by ModelFinder implemented in IQ-TREE [32]. RAxNG analysis included 200 non-parametric bootstraps, while IQTree incorporated 1000 ultra-fast bootstraps [37] with-allnni and-bnni settings.

The maximum likelihood (ML) trees (shown in Figure S2) are generally well supported (mean RAxNG non-parameteric bootstrap support 85% for Megabats and 85% Pigeons, with 75% and 78% respectively of nodes with BS>70). Both the Megabat and Pigeon trees are characterized by a long stem to the closest extant outgroup (stem to crown ratio >1) and a soft polytomy or basal radiation with short internodes; also seen in the phylogenomic trees [28,29] showing that it is not an artefact of insufficient data.

#### 2.3.1 Time tree estimation

We converted the RAxNG ML and 200 bootstrap trees into calibrated time-trees using Penalised Likelihood Rate Smoothing (PLRS) method [38], as implemented in treePL [39]. For the Megabats there are no reliable internal calibrations [40] but reasonably consistent extant crown age secondary calibrations from previous mammal studies [25,41,42]. Extant crown age root = 25 My, sd=2. Calibrations for pigeons follow previous studies [15,43]: secondary Columbimorphs (mean=58.0 My, sd=4.0), and two internal minimum fossil-based calibrations (*Rupephaps* and *Primophaps* [44,45]; but with *Primophaps* as stem ancestor).

TreePL was run on the full set of ML and bootstrap sample trees with log_penalty, ‘thorough prime’ optimized settings and smoothing factor drawn from a uniform distribution 5-20, and the secondary calibration drawn from a normal distribution (defined above). This provides a set of time-trees incorporating topological, branch length, calibration and relaxed-clock model uncertainties, which can then be summarized as a maximum clade credibility common ancestor tree in TreeAnnotator [46,47], analogous to Bayesian summary time-trees. These are shown in Figure S3. We chose PLRS because the shape of the trees (e.g. early radiation of short internodes) does not fit speciation models in Bayesian relaxed-clock methods (e.g BEAST), which also struggle with large amount of sequence data [24,48]. It also means the results reflect the more conservative non-parametric bootstrap topology support.

Dating resulted in similar late Oligocene ages for both bat and bird extant crown radiations (Figure S2). Megabats (25.1 mya 95%CI 21.2–29.0) merely returned the secondary calibration on the crown age. The Pigeon date of 26.8 mya (95%CI 25.1–28.6) is slightly younger than our previously published estimate (34.6 mya;[15]). This may reflect the increased power of the phylogenomic data combined with not enforcing a Bayesian speciation prior.

### 2.4 Historical biogeography modelling

To define a practical biogeographical categorization of regions, we balanced capturing the complexity of our focal region of the Indo-Australian Archipelago (IAA: the islands between Australia and continental south-east Asia), with limiting biogeographic model complexity. Megabats and Pigeons have similar concentrations of diversity and distribution across the IAA focal zone, hence Megabats regional assignment largely followed that of Pigeons [15]. The core IAA plus Asia regions are essentially the same for both taxa but megabats have an Africa region while pigeons have OldWorld (Europe plus Africa) and Americas (Table S5). Tables S1 and S2 include region assignments for each megabat and pigeon taxon.

We estimated ancestral states using two approaches: i) a Discrete-state Markov rate-based Maximum Likelihood (Mk-ML) model [49] run in the R package corHMM [50], and ii) a DEC model implemented in BioGeoBears [51]. The Mk-ML method used a simple single rate (ER) model with the Maddison and Fitzjohn (“maddfitz”) root probability. This is analogous to a DEC+J model. In our Megabat and Pigeon region schemes, 90% and 85% respectively of species are endemic to a single region, and the moderate proportion of multistate species is a tolerable level for the Mk-ML ancestral state method to accommodate.

#### 2.4.1 Summarizing biogeographic model inference

Rather than produce a single best ancestral state result, we focussed on broad biogeographical inferences summarizing both model ancestral state and phylogenetic uncertainty. To do this we applied stochastic node state mapping using 200 re-samplings of node state marginal likelihoods, to each of the PLRS-calibrated ML tree and 200 bootstrap trees. Transition events were summarized according to parent to daughter node (including tip) states; cases where the state remains the same were categorized as ‘endemic cladogenesis’. These results were divided up by divergence age into time-bins, assigned by daughter node age. For counting state lineages, branches spanning a time point were assigned to closest parent or daughter node state.

To investigate the role of islands in diversification and dispersal, we collated results into broader categories of continents (Americas, OldWorld/Africa, Asia, Australia) and islands (Wallacea, Melanesia, Philippines, Indian Ocean and Pacific Ocean) by post hoc combining states and state changes inferred in individual region analyses. We did not follow traditional conventions of extending Melanesia from New Guinea to Fiji, but instead including islands east of New Britain into ‘Pacific Ocean’. Summary results of events or states per time-bin are best interpreted as patterns of relative frequency or probability density rather than explicit counts. By summarizing the results across the whole marginal likelihood, the method accounts for multiple solutions and hence to some degree accounts for multistate solutions, particularly for combined region summaries. For phylogenetic visualization onto the single ML tree, we averaged marginal likelihoods across all 200 sample trees, for all nodes in common with the ML tree (identical taxon bipartitions) and the linked parent node [52]. Following O’Hara et al. [53], nodes (hence branches) with maximum state probability less than two thirds (<67%) were scored as indecisive.

#### 2.4.2 BioGeoBEARS

The probability of ancestral ranges was also estimated using BioGeoBEARS [51] in R (R Core Team, 2016) using the same trees and region distribution. We rationalized to a maximum of four regions and disallowed transitions between [Australia, Wallacea, Pacific Ocean, Philippines] and OldWorld/Africa. DEC+J was the best-fitting model for megabats, and DIVA+J for pigeons. Results were summarised as simplified circle trees with node marginal pie charts (Figure S5). Summaries of diversity and dispersal for the key categories (especially Australia) were essentially the same between Mk-ML and BioGeoBEARS. Differences in ancestral state inference for deeper nodes highlight how uncertain these are – to any method – due to the structure of the tree.

## 3. Results

### 3.1. Summary of phylogenetic results

For megabats the final phylogeny included 204 tips, comprising 171 species (84% of batnames.org v1.10 species) plus 11 subspecies and a further 22 extra Operational Taxonomic Units corresponding to populations. We rationalized sequences to additional subspecies/geographic units/distinct genetic units based on recognized nomenclature and coherent genetic data, particularly the extensive mitochondrial cytb, ND2 and COI data. This involved some re-interpretation of sample identities, resulting in taxonomic and phylogenetic novelties (see Supplementary file sections ‘Taxonomic and Nomenclature Notes’ and ‘Amendments to Nesi (2021) samples’). Combined with the full list of gene data information (see below) we present this as a contribution towards collating and describing available megabat genetic data. The updated pigeon phylogeny comprised 252 species (73% of IOC9.2 species), including all 22 species native to continental Australia.

Both the megabat and pigeon data resulted in generally well-supported trees (Figure S1) with a characteristic long stem and basal radiation, reflecting previous phylogenomic studies [28,29]. Dating resulted in similar late Oligocene ages for both bat and bird extant crown radiations (Figures 1, S2). For biogeographic inference megabats were coded into eight regions and pigeons into nine regions, with the extra region for the latter being the Americas (Table S5). For megabats regional diversity ranged from 41 (Pacific Islands) to 8 (Australia), and for pigeons from 60 (Americas) to 10 (Indian Ocean). Ancestral state biogeographic modelling by both Mk-ML and DEC-DIVA produced very similar results (Figure S5) so we concentrate on the summaries of the Mk-ML model integrated over 200 sample trees.

**Figure 1.**
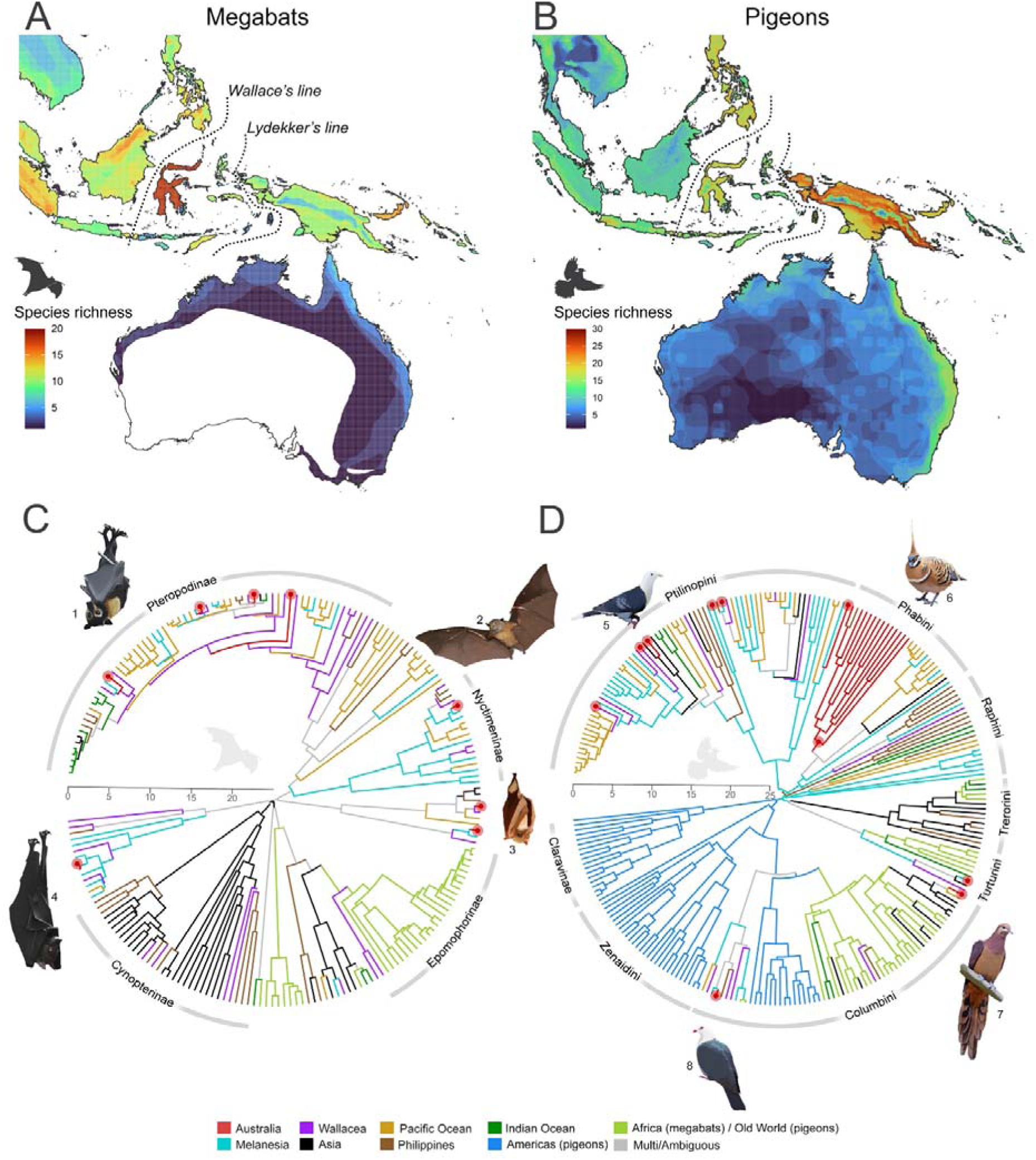
Indo-Australian Archipelago species richness map and global time-calibrated phylogenetic trees with ancestral areas reconstructions for Megabat (A and C, respectively) and Pigeon (B and D, respectively). Distributional data were extracted from the IUCN database and mapped onto a 10km² grid. Pigeon and Megabat chronograms show Mk-ML ancestral state reconstruction (common clade summary from 200 sample trees – see Methods; full trees shown in Figure S3). See Table S5 for region information. Grey indicates indecisive (maximum marginal <0.67). Timescale in millions of years. Region colour-coding used throughout. Illustrations: 1, *Pteropus conspicillatus*; 2, *Nyctimene robinsoni*; 3, *Macroglossus minimus*; 4, *Dobsonia moluccensis*; 5, *Ptilinopus alligator*; 6 *Geophasp plumifera*; 7, *Macropygia phasianella*; 8, *Columba leucomela*.

### 3.2. Early diversification

Megabats and pigeons both show a similar pattern of a long stem and a short basal radiation. In megabats Asia dominates this early radiation (71%), followed by Melanesia (14%) (Figures S6, S8), while for pigeons it is Americas (34%) and Melanesia (65%). Summaries of regional state lineage accumulation and endemic cladogenesis (Figures S6, S8) reinforce this pattern of early diversification in megabats concentrated to the west (Asia) but further east in Pigeons, particularly Melanesia.

### 3.3. Origins of Australian taxa

In megabats eight independent colonisations of Australia were inferred (Figures 1C, S3A, S8C), mostly from Melanesia, accounting for all species in Australia: the three endemics and populations of five species that also occur in nearby regions. Newly sampled Australian populations of *Macroglossus minimus* and *Synconycteris australis* show low divergence from Melanesian populations. The Australian endemic species *Nyctimene robinsoni* also shows very low mitochondrial divergence from other nominal species in southern New Guinea, indicating very recent geneflow between New Guinea and Australia (Figure S9).

In pigeons ten independent colonisations of Australia were inferred (Figures 1D, S3B, S8D), again dominated by Melanesia. The oldest colonisation involved a group of largely terrestrial and seed-eating species (the Phabines) that gave rise to near endemic radiation of 15 species (of which 13 occur in Australia). Of the remaining nine inferred colonisations of Australia, all involve primarily rainforest lineages, three led to species that are now endemic, while six represent populations of species that also occur in Melanesia or other nearby regions.

### 3.4. Regional comparisons of endemic cladogenesis and lineage accumulation

Summaries of regional state lineage accumulation and endemic cladogenesis indicate that megabats and pigeons have both been present and speciating within island arc systems (Philippines, Wallacea, Melanesia and Pacific Ocean islands) since start of the Miocene (Figures 2, S6 and S7). In both megabats and pigeons early island arc diversification is concentrated in Melanesia (Figures S6-8). Asia is the predominant centre of early megabat lineage accumulation and diversification, whereas Asian pigeon richness is generally low until the mid-Miocene (<15 mya).

**Figure 2.**
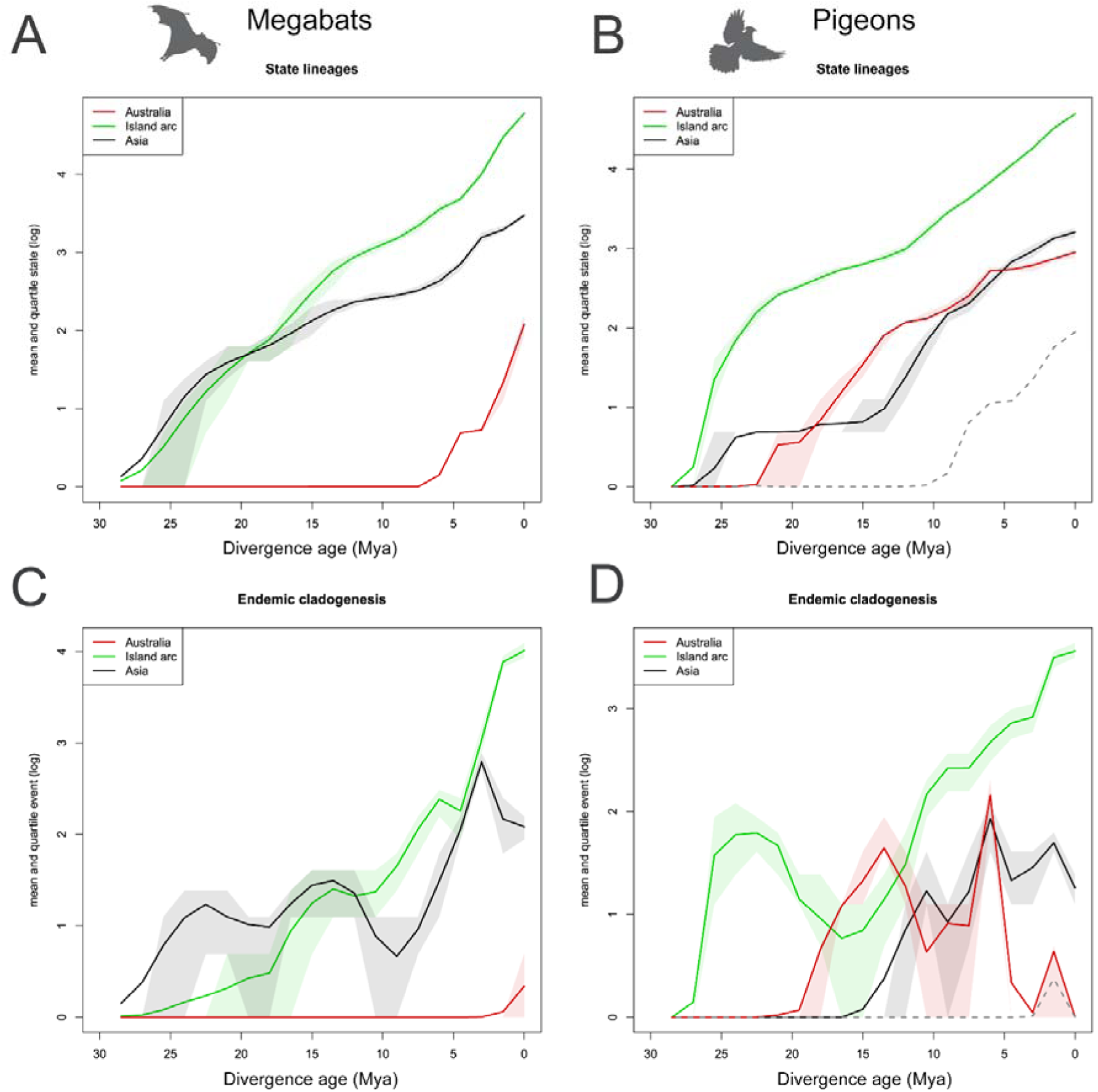
State Lineages and Endemic (within region) Cladogenesis for Megabat (A and C, respectively) versus Pigeon (B and D, respectively) differentiating between Island arc, Asia and Australia. Dotted line shows Australia without the Phabines. Note Asia includes India to Japan to Sunda. X-axis shows log of mean (line) and quartile range (shading) from 200 stochastic node mappings for 200 tree samples. Dotted line shows pigeon results discounting the Phabine clade.

Australia shows no signature of lineage diversity before the Plio-Pleistocene in megabats, and negligible endemic cladogenesis at all (Figure 2A, C). In pigeons Australian endemic cladogenesis and lineage accumulation is concentrated in the single radiation of largely terrestrial feeding and seed-eating Phabines (Figure 2B, D). Discounting this group, lineage accumulation only begins after the mid Miocene (<10 mya; with *Lopholaimus* being the oldest lineage), and the signature of endemic cladogenesis again is negligible (dotted lines in Figure 2B, D).

### 3.5. Upstream dispersal dynamics

In terms of upstream versus downstream dispersal processes (Figure 3A,B), both groups have similar numbers of inferred events but different temporal patterns (megabat: 20 upstream vs 27 downstream; pigeons: 22 upstream vs 21 downstream). Megabats show more of an older downstream pattern (consistent with Asia/Sunda early diversification) and more recent upstream. Pigeons however show both processes throughout their history. For pigeons Melanesia is the principal source of upstream dispersal (69%), while for megabats it is more evenly spread (33% Melanesia 22% Philippines 20% Wallacea).

**Figure 3.**
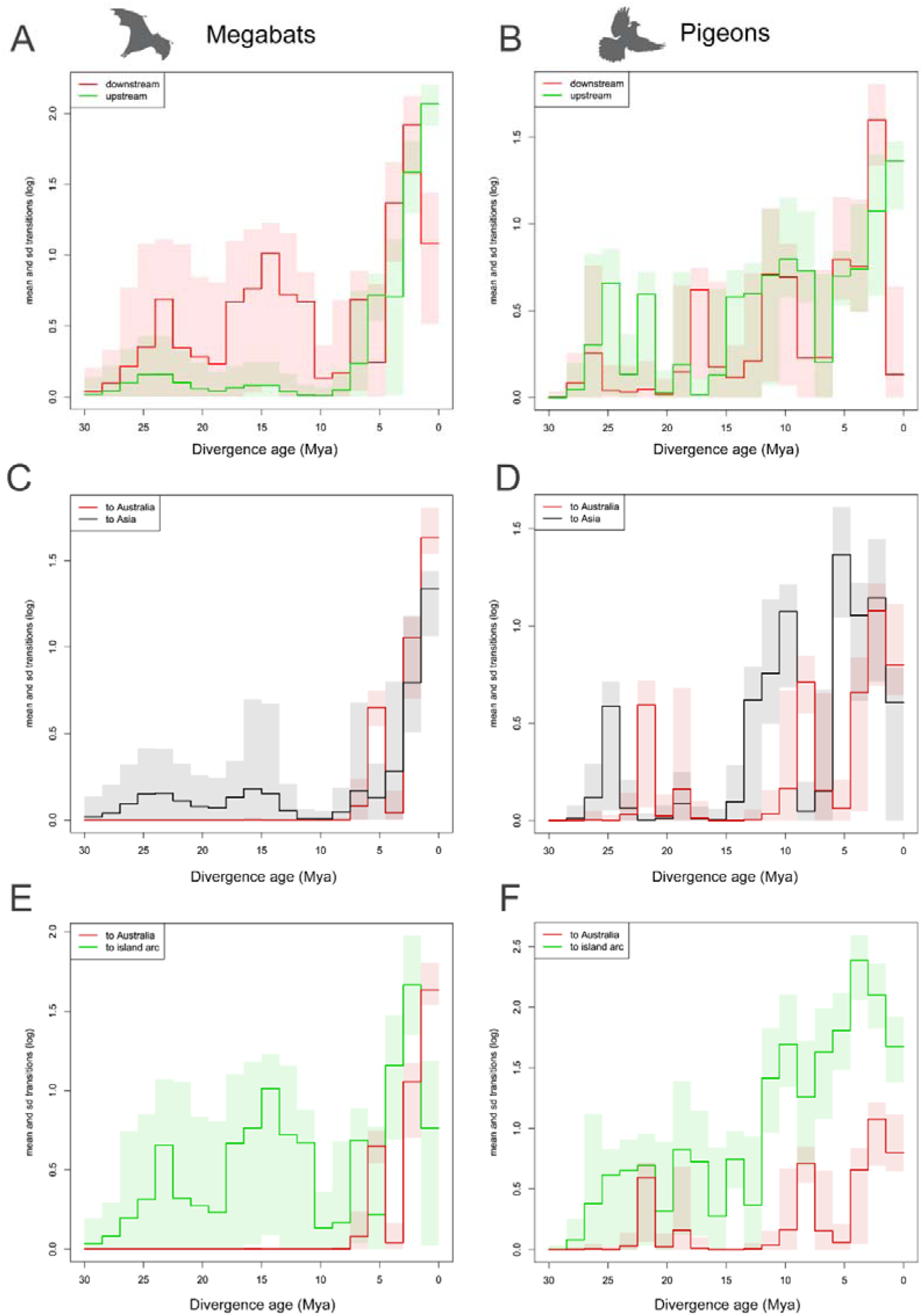
Comparisons between Megabat and Pigeon transitions within upstream/downstream (A and B, respectively), Australia/Asia (C and D) and Australia/Island arc (E and D). Upstream is all island to all continents, downstream vice versa. To island arc includes dispersals between island arcs as well as from outside into island arc. See Figure 1 and Table S5 for region information. X-axis as for Figure 2 but for transitions and with range (shading) based on ± one standard deviation.

Megabats and pigeons both share a marked increase in the number of inferred upstream colonisation rates into Australia over last 5 million years (Figure 3C, D). For megabats upstream colonisation into Asia is also largely only recent (Figure 3C). In pigeons, upstream dispersal into Asia and Australia has an older history, in part reflecting the early colonisation by different radiations (*Treron* in Asia, the Phabines in Australia) (Figure 3D). Across island arc regions (especially Wallacea, Melanesia, and Pacific Ocean islands) dispersal is inferred to be relatively ongoing process over the history of both groups, forming a striking contrast against comparatively low overall rates and marked temporal clustering of dispersals into nearby Australia (Figure 3E, F).

## 4. Discussion

Using the most species-complete phylogenetic datasets for megabats and pigeons assembled, we here show that Australian rainforest faunas in both radiations are dominated by lineages that arrived via numerous independent upstream colonisations during the Plio-Pleistocene. While upstream colonisation of continents from islands has now been documented in an array of taxa and different parts of the world [3], such a temporally concentrated dominance of upstream colonisations in the assembly of a continental community is novel. This finding has important implications for both our understanding of the history of Australia’s rainforests, and for the role that upstream colonisation may play in shaping regional biotas.

In total we estimate a minimum of 17 colonisations of Australia from islands by forest-associated taxa (eight bats, nine pigeons). The true number of dispersals is likely higher as it seems likely that different populations within widespread and mobile species in genera such as *Ducula*, *Macroglossus*, *Pteropus* and *Ptilionopus* may have colonised different parts of northern Australia independently. These upstream colonists also show little evidence for subsequent endemic cladogenesis within Australia, indicating that dispersal is the source of all new taxonomic diversity. For megabats, this biogeographic inference spanning all genera is novel, and provides an output consistent with the dearth of fossil megabats in Australia, even in mammal-rich fossil assemblages where other lineages of (microchiropteran) bats are abundant [54]. Seven other bat families have been found in Australian fossil deposits dating back to the Miocene, while the oldest megabat fossil in Australia is late Pleistocene [54]. For pigeons our inference from an expanded and species complete sampling of Australian taxa and revised dating strengthens previous inferences [15].

In contrast to the overall recency of the Australian megabat and rainforest pigeon assemblages, syntheses presented here and previously published analyses [28,55,56] all also show that islands to the north are characterised by more ecologically diverse, species-rich and older clades. Early pigeon diversification (and also contemporary alpha diversity) is concentrated towards Melanesia and the Philippines, perhaps reflecting regional origins via westwards colonisation from South America. Megabat diversity is more concentrated towards Sunda-Wallacea, likely reflecting a deeper history and potential origins of this group in continental Asia. Despite these differences, biogeographic inferences for both taxa support the broader hypothesis that island systems have been subaerial and generating biodiversity across Australasia since before the Miocene [57,58]. Dispersals between islands and continental Asia have also overall occurred more frequently than between islands and Australia (Figure 3). Indeed, two megabat genera in the Sunda-Wallacean region (*Chironax and Aethalops*) appear deeply divergent and paraphyletic, further emphasising the deep history across Asia-Wallacea and contrasting with the overall recency of the continental Australian fauna.

Our findings here contrast with an entrenched understanding of Australian rainforest biotas as dominated by relictual taxa that have retreated into tiny islands of refugial habitat in response to intensifying aridification [10]. Given the dominant role that megabats and fruit pigeons play in long-distance dispersal and pollination in contemporary Australian rainforests [20,21], the recency of these keystone faunas is particularly notable. It seems likely that the mass influx of volant herbivores into Australia documented here, may have also driven concomitant change in the floral composition of lowland tropical habitats in Australia. In support of this idea, synthetic analyses of the biogeographic origins of Australian rainforest plants and birds that have found a similar – if less marked - dominance of “immigrant” lineages with tropical origins in megathermal habitats in eastern and northern Australia [16,19]. Australian rainforest plant taxa with immigrant origins also tend to have smaller fruits than those with deeper “Gondwanan” histories, a pattern consistent with a dominant role for volant herbivores in their recent dispersal [17]. We hypothesise that with adequate sampling from New Guinea and with appropriate biogeographic coding, future analyses of rainforest tree radiations may reveal that at least some may show an overlooked signature of concerted recent upstream colonisations into Australia from New Guinea to match that we detect in volant herbivores. One potential example comes from phylogenetic analyses of a major clade within one of the most important lineages of rainforest trees (*Ficus*) which provided evidence of extensive diversification in Melanesia, contrasting against an Australian biota mostly comprised of populations of widely distributed species [59]. We also hypothesise that increasing spatial overlap with volant herbivores may have played a role in the rapid radiation and spread of some rainforest tree genera with Australian origins, especially the mega-diverse genus *Syzigium* [60].

With respect to understanding why megabats and pigeons show such high rates of upstream colonisations into Australia, the key questions are what factors prevented successful colonisations, and what then changed to open the way through the Plio-Pleistocene? Alternative explanations could invoke either geographic, environmental and biotic barriers impeding successful colonisation until recently.

Both megabats and pigeons include many taxa that are excellent dispersers capable of traversing over hundreds or more kilometres of open ocean [61,62]. Furthermore, while the history of land connection between Australia and New Guinea through the Miocene is complex, both phylogenetic and geological evidence indicate at least periodic land connections going back at least as far as the mid-to late-Miocene [11,63]. In this context geographic barriers seems a limited explanation for low rates of dispersal into Australia until recently.

It seems more plausible that through the late Tertiary, the temperate and more strongly seasonal environments that still dominate much of Australia may have been difficult for largely tropical forest lineages to colonise. In support of this idea, in contemporary Melanesia where megabats and rainforest pigeons show centres of diversity and have deep history, species richness is comparatively low in cooler habitats at higher elevations (Figure 1), and in the case of megabats dominated by nectar specialists [64]. Some environmental reconstructions suggest temperate environments may have been much more widespread in the past in Australia when the continent was further south, and that tropical environments were limited or even absent [65]. Two of the most divergent megabat and pigeon lineages present in Australia (*Pteropus scapulatus* and *Lopholaimus antarcticus*) are also both particularly nomadic and follow food resources huge distances [61,62], further indicating that irregular and seasonal Australian environments may have been key constraints on establishment, and then perhaps drivers of evolution.

An alternative environmental barrier could be differing biotic communities between regions preventing establishment, and it has even been suggested that widespread extinction of arboreal marsupial herbivores may have opened the way for megabat colonisation over the late Tertiary [66]. This hypothesis is difficult to test. However, given the shared upstream signature across diurnal pigeons and nocturnal megabats, and the success and diversity of megabats in herbivorous mammal-rich communities in south-east Asia and Africa (Figure 1), we tend to favour the climatic barrier hypothesis.

If climatic and environmental shifts within Australia have been the primary factor responsible for the recent upstream influx of volant forest herbivores, then a further implication is that the manifest pattern is in many respects consistent with core predictions of island biogeography [1]. In short, the region with largest area and oldest lowland tropical forest (islands, and especially New Guinea) has been functioning as a source area for restricted and putatively young lowland tropical habitats in eastern and northern Australia. A more global analysis of the extent to which upstream colonisations are associated with disturbed, marginal and/or relatively young environments could add important nuance to our understanding of the role of this process in shaping regional biotas. We hypothesise that such an analysis could reveal that when factors such as habitat type, area and species richness are considered, many upstream colonisations may not be anomalous with respect to underlying predictions of island biogeography. For instance, a number of studies showing upstream colonisations into south-east Asia have argued that these dispersals have occurred into lower biodiversity habitats such as mangroves or montane areas [15,58]. Building on these idiosyncratic examples, we here demonstrate a case wherein rich insular communities of excellent dispersers have served as a dominant diversity source for rainforest assemblages on a nearby and climatically unstable continent.

## Data accessibility statement

Supplementary files are available upon request to the authors, pending publication by the journal.

## Acknowledgments

We thank Julien Louys and Sue Hand for advice about the fossil record for Australian bats; and Janne Torkkola for work on additional sequencing. We also acknowledge the numerous researchers who have provided the specimens and genetic data that has underpinned this work.

